# Genome-wide divergence among invasive populations of *Aedes aegypti* in California

**DOI:** 10.1101/166629

**Authors:** Yoosook Lee, Hanno Schmidt, Travis C. Collier, William R. Conner, Mark J. Hanemaaijer, Montgomery Slatkin, John M. Marshall, Joanna C. Chiu, Chelsea T. Smartt, Gregory C. Lanzaro, F. Steve Mulligan, Anthony J. Cornel

## Abstract

In the summer of 2013, *Aedes aegypti* Linnaeus was first detected in three cities in central California (Clovis, Madera and Menlo Park). It has now been detected in multiple locations in central and southern CA as far south as San Diego and Imperial Counties. A number of published reports suggest that CA populations have been established from multiple independent introductions. Here we report the first population genomics analyses of *Ae. aegypti* based on individual, field collected whole genome sequences. We analyzed 46 *Ae. aegypti* genomes to establish genetic relationships among populations from sites in California, Florida and South Africa. We identified 3 major genetic clusters within California; one that includes all sample sites in the southern part of the state (South of Tehachapi mountain range) plus the town of Exeter in central California and two additional clusters in central California. A lack of concordance between mitochondrial and nuclear genealogies suggests that the three founding populations were polymorphic for two main mitochondrial haplotypes prior to being introduced to California. One of these has been lost in the Clovis populations, possibly by a founder effect. Genome-wide comparisons indicate extensive differentiation between genetic clusters. Our observations support recent introductions of *Ae. aegypti* into California from multiple, genetically diverged source populations. Our data reveal signs of hybridization among diverged populations within CA. Genetic markers identified in this study will be of great value in pursuing classical population genetic studies which require larger sample sizes.

## Introduction

*Aedes aegypti* has a short flight range, usually not actively moving more than 200 m from their breeding source ^1^, but is exquisitely adapted to hitchhiking in transport vehicles ^2^. One central question concerning the populations dynamics of *Ae. aegypti* in CA therefore is whether the established populations at different locations are founded from one source population that spread across the state or if they are the result of other kinds of founding effects. A recent study revealed several, genetically distinct *Ae. aegypti* populations in CA presumably originating from multiple introductions from other sites in the U.S. and/or northern Mexico ^3^. Insight into the population structure of CA *Ae. aegypti* beyond this will be necessary to fully understand the dynamics that shape the current pattern of distribution of this invasive vector species.

California (CA) had no known established local populations of *Ae. aegypti* prior to the summer of 2013 when it was detected in three cities in central California: Clovis, Madera and Menlo Park ^4, 5, 6^. In the spring and summer of the following year, this mosquito was again found in the same three California locations and for the first time in additional communities in central California and further south in San Diego County (Figure 1). *Ae. aegypti* specimens have been collected over multiple years from some sites and from additional locations in both central and southern CA each year, indicating that *Ae. aegypti* has now become established and is spreading through large parts of the state (Figure 1).

**Figure 1.**
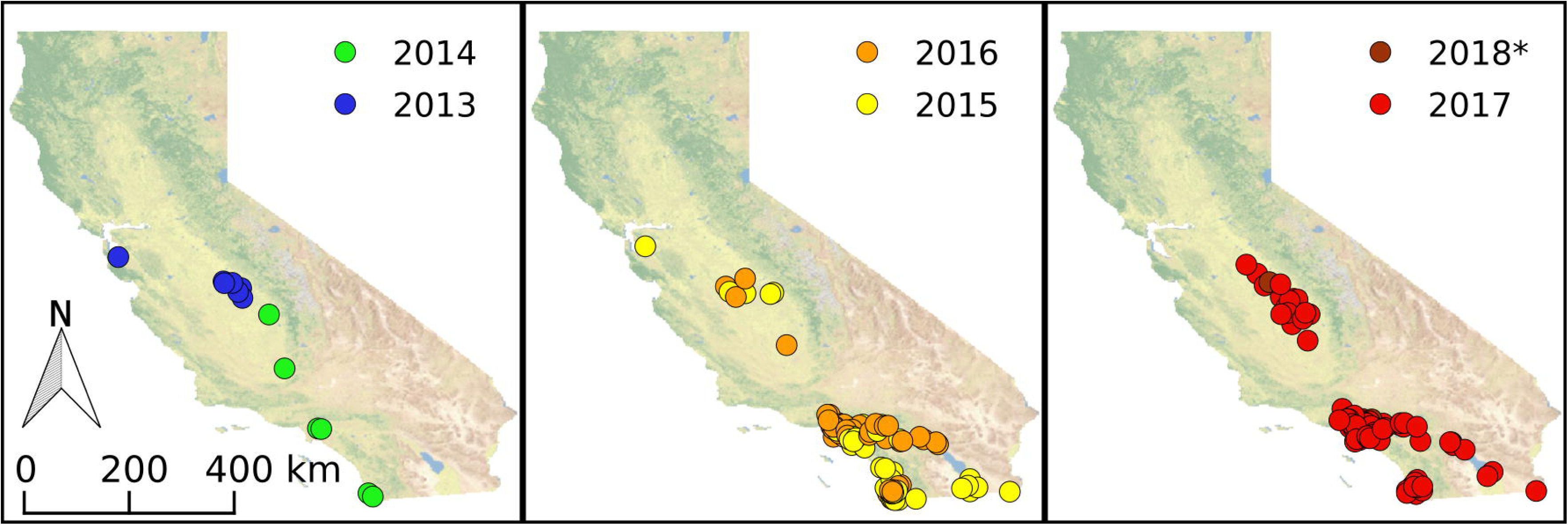
History of the recent *Ae. aegypti* invasion in California. The respective year of the first detection of the species in each site is displayed on the map. * 2018 data is surveillance data available as of July, 31, 2018. Data was derived from CalSurv Maps ^38^ and the CleanTOPO2 basemap ^39^ was used as background.

*Aedes aegypti* is thought to have a short flight range, usually not actively moving more than 200 m from their breeding source ^1^, but is exquisitely adapted to hitchhiking in transport vehicles ^2^. Recent studies revealed several, genetically distinct *Ae. aegypti* populations in CA originating from multiple introductions from other sites in the U.S. and/or northern Mexico ^3^.

Successful vector control can benefit from population genetics and genomics analyses which can provide estimates of gene flow and identify the genetic basis of phenotypes such as insecticide resistance ^7^ and host preference ^8^. Population genomics studies are especially critical to the development of control strategies based on genetic manipulation of vectors, which is a matter of growing interest. Modelling, planning and monitoring activities associated with control programs require affordable and rapid assays to distinguish vector sub-populations within a species and a deep understanding of the processes that shape their genetic structure. It is becoming increasingly apparent that hybridization between diverged vector populations may be an import source of new genetic material including alleles that mediate adaptations to facilitate range expansion ^9^ or that promote the evolution of resistance to insecticides ^10^. Analysis of whole-genome sequencing data is the most powerful method to detect even minor admixture ^11^. Therefore, we have applied a population genomics approach to study invasive *Ae. aegypti* populations. Here we report a preliminary analysis based on genome sequences of 39 individual *Ae. aegypti*, collected from twelve locations throughout CA, and, for comparison, four specimens from Florida and three from South Africa. The analyses presented here should serve as a starting point for expanded population genomics studies aimed at understanding how invasive mosquito species become established in new locations and how distinct populations interact on the genetic level.

## Materials and Methods

### Mosquito Collection

Adult female *Ae. aegypti* were collected from 13 cities by personnel from Mosquito Abatement Districts in Fresno, San Diego, and Orange Counties (Figure 2 and Supplementary Table S1). All collections on private properties were conducted after obtaining permission from residents and/or owners. Mosquito samples were individually stored in 80% ethanol prior to DNA extraction.

**Figure 2.**
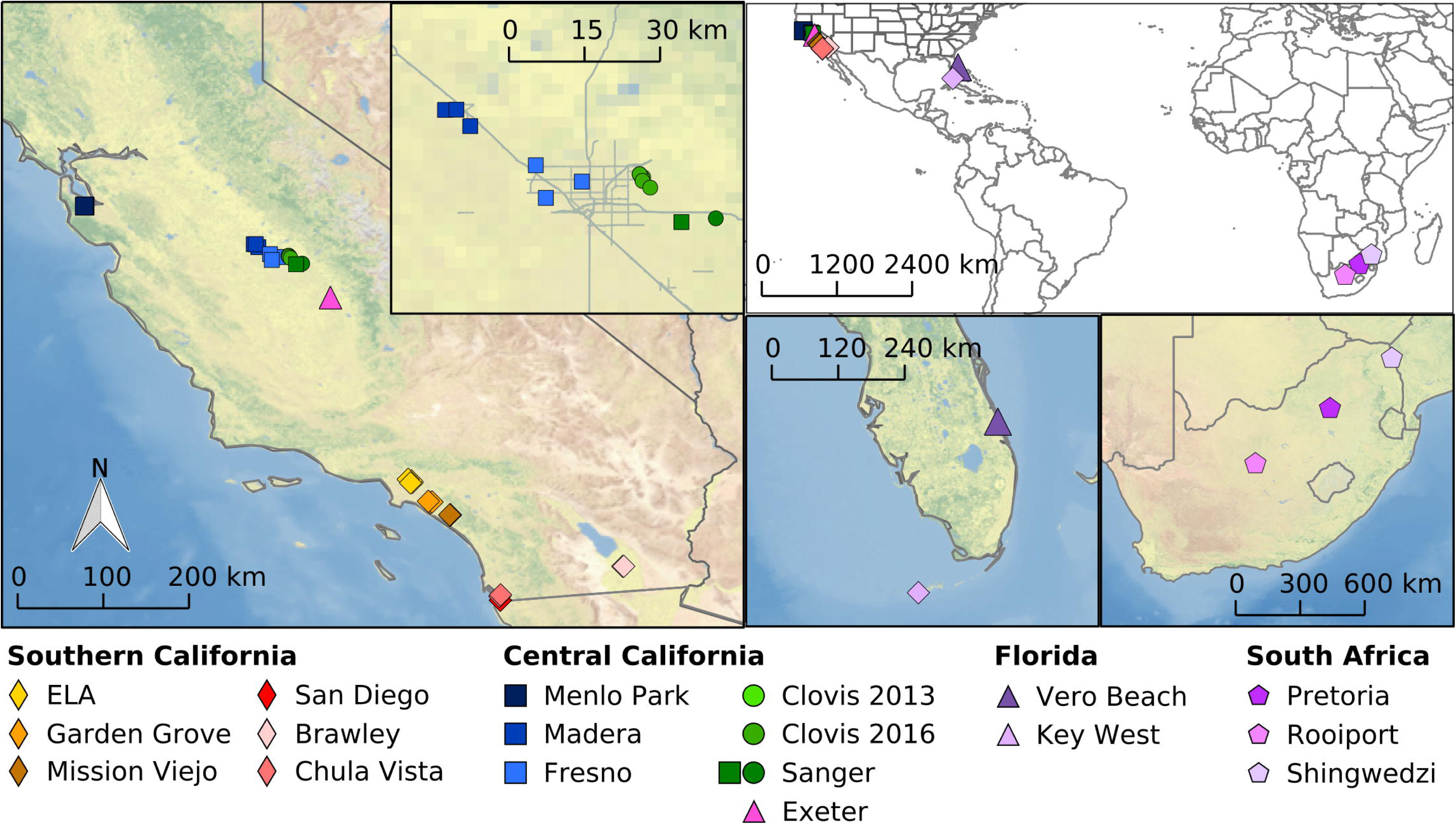
Geographic origin of *Ae. aegypti* used in this study. Left map shows location of all samples from California with the inset enlarging the Fresno/Clovis area. Top right shows a world map, bottom central shows a Florida map and bottom right a South Africa map with all respective sampling locations. CleanTOPO2 basemap ^39^ was used as background.

### Whole genome sequencing

Genomic DNA was extracted using established protocols ^12, 13^. DNA concentrations for each sample were measured using the Qubit dsDNA HS Assay Kit (Life Technologies) on a Qubit instrument (Life Technologies). A genomic DNA library was constructed for each individual mosquito using 20 ng DNA, Qiaseq FX 96 (Qiagen, Valencia, CA), and Ampure SPRI beads (Beckman) following an established protocol ^13^. Library concentrations were measured using Qubit (Life Technologies) as described above. Libraries were sequenced as 150 bp paired-end reads using a HiSeq 4000 instrument (Illumina) at the UC Davis DNA Technologies Core.

### Sequence Analysis

Raw reads were trimmed using Trimmomatic ^14^ version 0.36 and mapped to the AaegL5 reference genome ^15^ using BWA-MEM ^16^ version 0.7.15. Mapping statistics were calculated using Qualimap version 2.2 ^17^(Table S1). Freebayes ^18^ version 1.0.1 was used for variant calling without population prior. We used minimum depth of 8 to call variants for each individual ^19^. Only biallelic SNPs were used for further analysis. A missing data threshold of 20% was used to filter SNPs. A phylogenetic tree base on the polymorphism data was constructed using the neighbor-joining algorithm as implemented in PHYLIP ^20^ version 3.696. Hudson F_ST_ ^21^, nucleotide diversity (π) and Principal Component Analysis (PCA) analyses was done in Python version 3.4.3 using the scikit-allel module version 0.21.1 ^22^. The presence of mitochondrial pseudogenes in the nuclear genomes of *Ae. aegypti* could potentially confound SNP calling ^23^. Thus we followed the mapping recommendations suggested by Schmidt, et al. ^24^ and mapped raw reads to the mitochondrial reference genome prior to mapping unmapped reads to the nuclear genome.

We used Ae13CLOV028MT (Genbank ID: MH348176) as a reference for mapping the mitochondrial genome because all our specimens contained a deletion between position 14,522 and 14,659 compared to the AaegL5 reference genome ^24^. Variants in the mitochondrial genome were called with Freebayes as described for the nuclear genome, but set to single ploidy. Mitochondrial coverage was on average 160 times greater than the nuclear genome coverage with a minimum of 25-fold difference (Supplementary Table S1). Use of properly paired reads for variant calling reduced errors generated by failing to recognize mitochondrial pseudogenes present in the nuclear genome. The Vcf2fasta program ^25^ was used to extract mitogenome sequences from the VCF file to FASTA format. MEGA version 7.0.26 ^26^ was used for mitogenome alignment. Mitogenome reference sequences of Culex quinquefasciatus (Genbank accession number = HQ724617), Aedes notoscriptus (KM676219), and *Aedes albopictus* (NC_006817) were obtained from GenBank and added to the alignment. Sequences for the thirteen mitochondrial protein-coding genes in *Ae. aegypti* were obtained from GenBank ^27^, extracted from our dataset, and concatenated for tree construction with the maximum likelihood algorithm implemented in MEGA.

### Data Visualization

QGIS version 2.18 was used to create maps. Python matplotlib version 2.2.2 (https://matplotlib.org/) was used for generating plots. Inkscape (https://inkscape.org/) version 0.92 was used to edit images.

## Results

We sequenced the genomes of 46 specimens of *Ae. aegypti* from California (N=39), Florida (N=4) and South Africa (N=3) with median genome coverage of 9.6X per sample (Supplementary Table S1). Sequence data is available through NCBI Sequence Read Archive (Study accession number: SRP106694). Data is also available through UC Davis PopI OpenProjects AedesGenomes page ^28^: Filtering for biallelic SNPs and a minimum depth of 8 with at most 20% missing data yielded 2,919,917 final high-quality SNPs.

Principal Components Analysis based on the genotypes of these SNPs revealed four distinct genetic clusters (Figure 3). The genetic cluster designated GC1 includes all six cities in southern CA and includes one central CA site, Exeter. Samples from Florida also fall within this cluster. The GC2 cluster includes the northernmost sites near the coast in Menlo Park and includes some central CA populations at Madera and Fresno. Populations from the restricted area around Clovis and Sanger in central CA form the GC3 cluster. The GC4 cluster includes all South African *Ae. aegypti* samples. Overall, the distribution of the three *Ae. aegypti* genetic clusters containing CA *Ae. aegypti* have a nearly parapatric distribution with the three groups potentially converging in the Central Valley. The population at Sanger appears to have multiple genetic clusters occurring in sympatry. One specimen from Sanger (Ae17CON058) could be GC2 or a hybrid of GC2 and GC3 given its values of PC2 and PC3 relative to other samples (Figure 3). Other Principal components (PC5 and PC6 in Figure 3) indicate additional population subdivisions further dividing Mission Viejo, Garden Grove, Exeter and Vero Beach samples from the rest of GC2 (Figure 3). This is consistent with a previous report suggesting highly structured populations within southern CA *Ae. aegypti* ^3^. Our genome-wide SNP-based clustering results show clear subdivision between Clovis and Fresno/Madera/Menlo Park. This may seem slightly different from published SNPchip data ^3^ but is consistent with the microsatellite-based genetic clustering reported in the same study.

**Figure 3.**
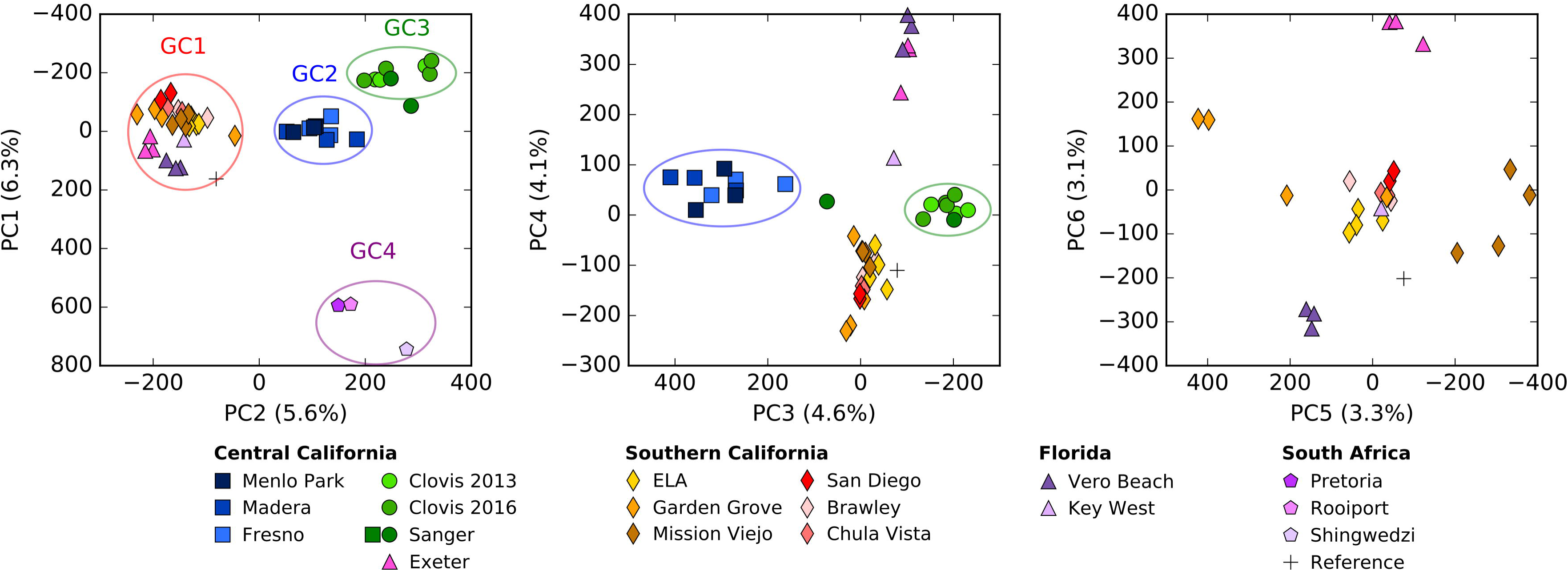
Genetic Clusters of *Ae. aegypti* based on PCA analysis. Principal Component Analysis based on the SNP data. Principal component 1 (PC1) accounts for 6.3% of SNPs that are separating South Africa population (GC4) from United States (US) populations. PC2 accounts for 5.6% of SNPs that separates US populations into three groups (namely GC1, GC2 and GC3). PC3 accounts for 4.4% of SNPs separating Central California population into two groups (GC2 and GC3). PC4 accounts for 4.2% of SNPs separating Southern California population from Exeter (Central California) and Vero Beach, Florida populations. PC5 (3.2% SNPs) and PC6 (3.1% SNPs) further divide GC1 cluster separating Exeter, Garden Grove and Mission Viejo and Vero Beach samples.

Windowed-F_ST_ analysis indicates genome-wide differentiation among the four genetic clusters (Figure 4A-C). The GC1 and GC4 clusters are the most highly diverged (genome-wide average F_ST_ = 0.159). Genetic distance between GC1 and the other clusters is intermediate (Figure 4 A and B). Nucleotide diversity (π) is highest in the middle region of each chromosome (Figure 4). These regions correspond to the location of the centromeres (coordinates obtained from personal communications with M. Sharakhova at Virginia Polytechnic Institute). Overall nucleotide diversity is lowest on Chromosome 1, which contains the sex determining locus in *Ae. aegypti*. Overall nucleotide diversity is similar in all population comparisons as indicated by difference in nucleotide diversity (Δπ = π_1_ – π_2_). As a peculiarity, the South African populations have noticeably higher nucleotide diversity in a region around 160-170 Mbp of Chromosome 1 compared to samples from the southern CA GC1.

**Figure 4.**
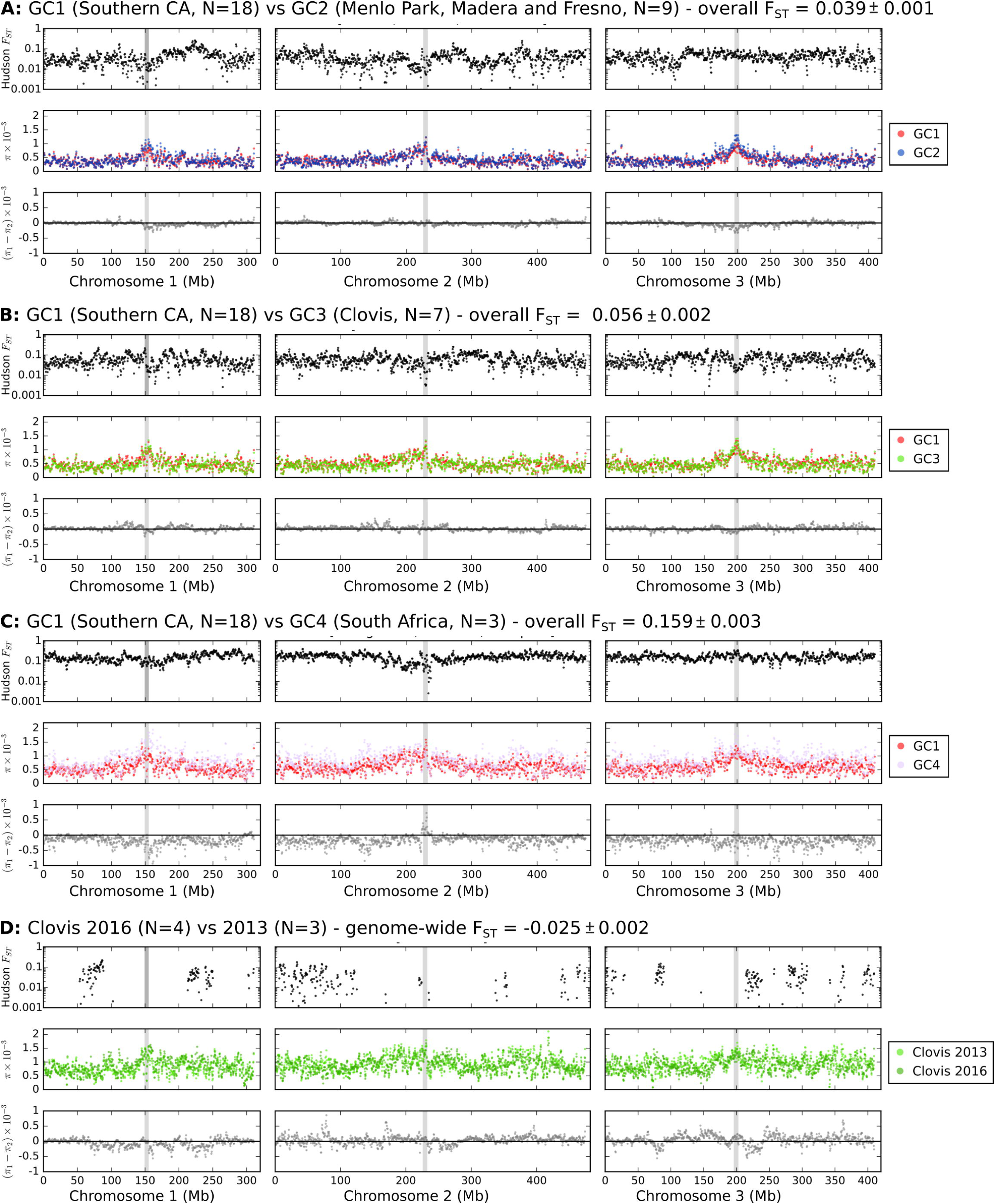
Genome-wide comparison of *Ae. aegypti* populations revealing broad highly differentiated (F_ST_>0.1) genomic regions. F_ST_ is marked in black dots. Nucleotide diversity (π) is marked in red for GC1, blue for GC2, green for GC3 and light purple for GC4. Difference in nucleotide diversity (=π_1_-π_2_) is marked in in gray. A: GC1 (Southern California, N=18) vs GC2 (Central California – Menlo Park, Madera and Fresno, N=9). B: GC1 (Southern California, N=18) vs GC3 (Central California – Clovis, N=7). C: GC1 (Southern California, N=18) vs GC4 (South Africa, N=3). D: Clovis 2013 (N=3) vs 2016 (N=4). Values were calculated using 1Mbp windows with 500Kbp steps. Vertical gray bars indicate the location of centromeres. Other comparisons of GC2 vs GC3, GC2 vs GC4 and GC3 vs GC4 are provided in Supplementary Figure S1.

For our only temporal comparison, we compared the genomes of samples obtained in Clovis in 2013 with those from samples collected in 2016. Overall genome divergence is negligible (F_ST_ = −0.025 ± 0.002). However, a whole genome scan using 1 Mbp windows for F_ST_ values indicates a number of genomic regions with markedly elevated F_ST_ values (>0.1) (Figure 4D). The difference in nucleotide diversity between 2013 and 2016 samples shows a reduction in nucleotide diversity over time in chromosome 2 and 3 (mean π_2013_ - π_2016_ value of 4.48×10^−5^ and 5.20 ×10^−5^, respectively). On the contrary, the nucleotide diversity increased over time on Chromsome 1 (π_2013_ - π_2016_ value of −6.36×10^−5^). However, an increase in nucleotide diversity in 2016 compared to 2013 is visible in all three chromosomes, some of which also coincides with highly differentiated (F_ST_ >0.1) regions. These highly differentiated regions with relative large nucleotide diversity change may indicate genomic regions under selection presumably as the founding population adapts to local environmental conditions ^7^.

Differentiation among populations within GC2 (F_ST_ =-0.052 ˜ −0.007) and GC3 (F_ST_ =-0.131 ˜ - 0.045) is minimal (Table S2). However, differentiation between some populations within GC1 is fairly high (F_ST_ =0.006 ˜ 0.210), especially between Exeter and the southern CA samples (F_ST_=0.094 ˜ 0.197). This suggests potential substructure within GC2 separating Exeter and Florida samples from other southern CA samples (Figure 3 and Figure 5). The F_ST_ distance between one sample from the town of Sanger (sample Ae17CON058) to GC2 and GC3 clusters were equivalent (Table S2) and its placement in the PCA (Figure 3) and phylogenetic tree (Figure 5) suggest that this individual could be a GC2/GC3 hybrid.

**Figure 5.**
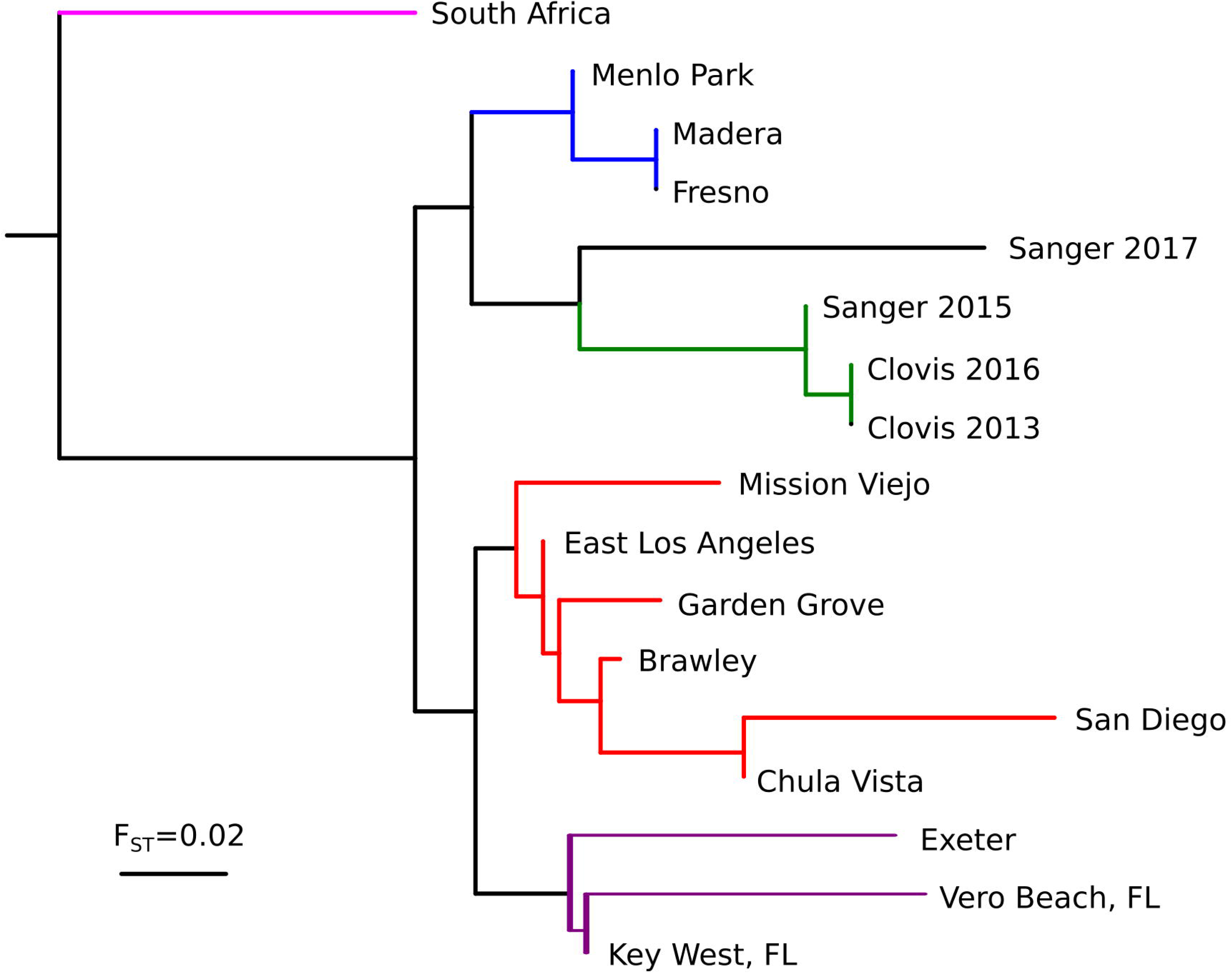
Neighbor-joining tree based on pairwise nuclear genome-wide F_ST_. Populations within GC1 designated in red (CA) and purple (Florida and Exeter, CA), GC2 in blue, GC3 in green, and GC4 in magenta.

Estimates of F_ST_ derived from whole genome sequence data have been shown to be accurate even with very small sample sizes (i.e. N=2/population ^29^). This is due to the very large number of SNPs (i.e. n>>1,000 loci) used in these analyses. Using conservative SNP calling with a minimum depth of 8 we were able to calculate F_ST_ values between pairs of genetic clusters based on 2.1 – 6.3 million SNPs (mean = 4.0 million). This well exceeds the number of loci required for an accurate assessment of F_ST_. In addition, we evaluated various read depths and missing data ratios and observed results consistent with those we report here (data not shown).

Mitogenome sequence analysis revealed two major mitochondrial lineages in CA *Ae. aegypti* (Figure 6). One lineage includes the *Ae. aegypti* reference sequence and is represented in all four genetic clusters, the other was present in samples from GC1, GC2 and GC4. Samples from Florida and South Africa are distributed among the two major lineages suggesting that these lineages might be present throughout the global range of *Ae. aegypti*.

**Figure 6.**
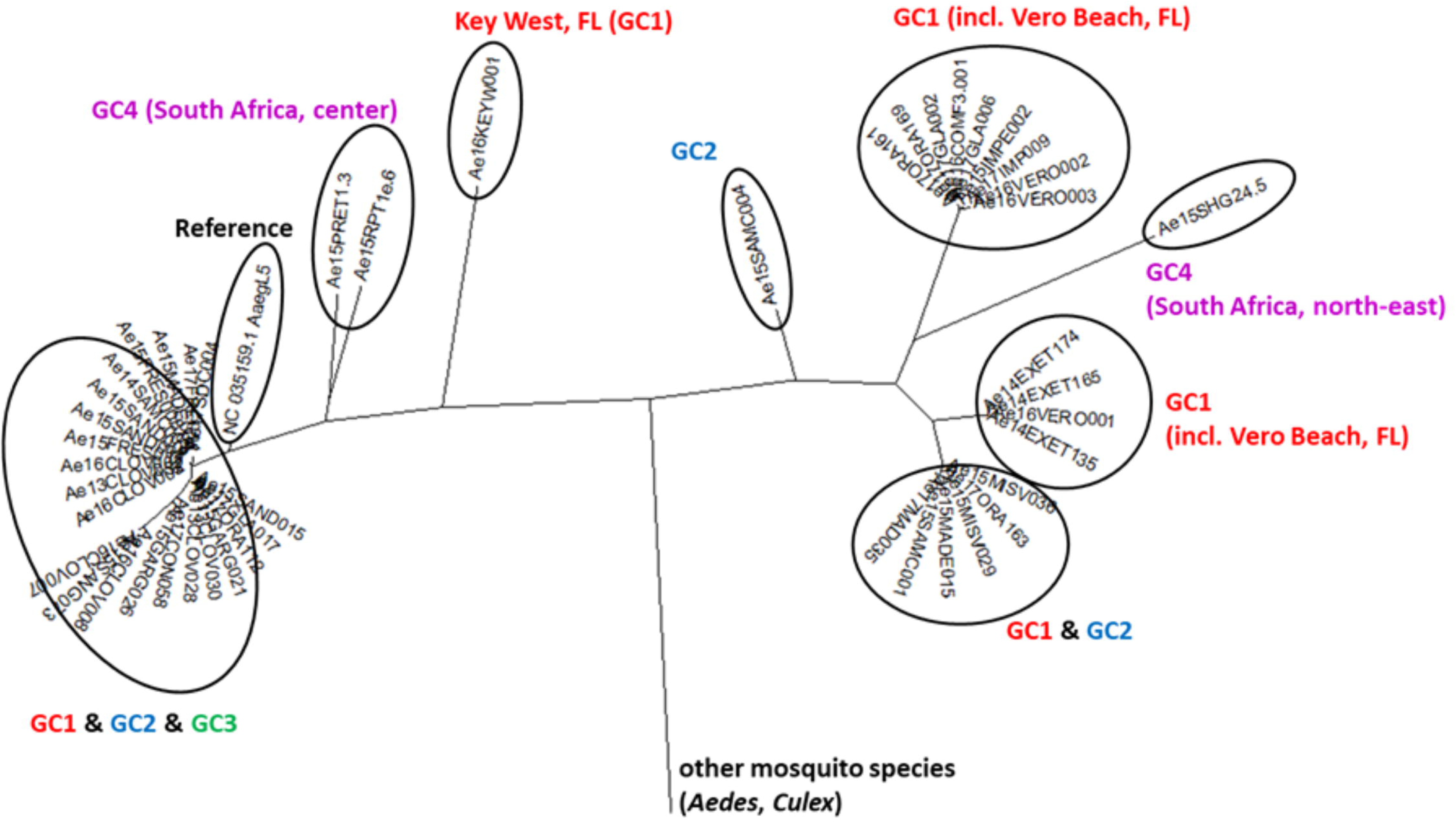
Maximum Likelihood tree based on mitochondrial protein-coding genes. Sample-specific sequences for all protein-coding genes were aligned and concatenated. Labelling refers to the Principal Component Analysis and resulting genetic clusters (GCs).

## Discussion

Using nuclear genome sequence data we identified three major genetic clusters among CA *Ae. aegypti*. These correspond roughly to geographic regions in the state (Figure 2 and 3). Our data support the hypothesis that *Ae. aegypti* in CA currently exists as multiple, mostly isolated populations. High genetic distance (F_ST_ > 0.1) as well as genome-wide differentiation (Figure 4) support multiple introductions into CA from genetically distinct source populations as the most plausible history of this invasion.

Two major mitochondrial lineages are present within California populations, probably corresponding to previously described global clades ^24, 30^. However, their genealogy differs from the nuclear genome genealogy (Figures 5 and 6). This is comparable to a previous study using ND4 sequence analysis of *Ae. aegypti* populations introduced to Florida ^31^. The lack of geographic clustering of mitochondrial lineages therefore appears to be common in invasive *Ae. aegypti* populations and is likely due to the saltatory nature of dispersal in this species. The incongruence between nuclear and mitochondrial gene genealogies could be due to different evolutionary rates between different loci producing differing topologies ^32, 33, 34^. It is possible that mitochondrial lineages capture historic divergence events, while nuclear genome divergence reflects relatively recent divergence. Linkage disequilibrium decays rapidly in mosquito genomes as seen in Anopheles arabiensis ^35^. Thus, any contact between two distinct *Ae. aegypti* populations may have resulted in relatively recent gene flow homogenizing populations within a locality. In this case mitochondrial markers appear to be less useful to determine relatively recent population divergence events in *Ae. aegypti*.

We investigated the possibility of generating SNP genotypes that are compatible with the existing *Ae. aegypti* SNP chip dataset ^36^ to allow for a direct comparison of our results with those previously published. The SNP positions provided in Evans et al. ^36^ were based on the initial genome assembly AaegL1 ^37^. Our BLAST results comparing AaegL1-based SNP sequences to the AaegL5 assembly revealed numerous and significant differences, including duplicated regions (multiple matching with high (>98%) similarity), sequence differences (arising from indel mutations), non-biallelic SNPs, polymorphisms surrounding the target SNPs, etc. These often resulted in mismatched genotype calls between the two different platforms, most likely displaying at least partially unintended haplotype calls with the SNP chip. Due to these problems a direct comparison of SNP genotype calls using the published SNP chip data with those generated from genome sequence data is deemed inappropriate and we highly recommend taking this into account when applying SNP chip analyses in the future.

Microsatellite data from Gloria-Soria, et al. ^4^ and Pless, et al. ^3^ indicated that San Mateo (=Menlo Park), Madera and Fresno samples were genetically similar to samples from the southeastern USA which includes samples from Louisiana, Georgia and Florida. Pless, et al. ^3^ also included a population from Exeter, CA that was also classified together with other central CA samples and south central and southeast USA populations based on microsatellite profiles. This appears to be inconsistent with our results. Our analysis placed the three central CA populations (Menlo Park, Madera and Fresno) in a group (GC3) distinct from the group containing the southeast USA populations (Vero Beach and Key West, Florida). In our analysis, the samples from Florida clustered with populations from Exeter, CA and southern CA (GC1, Figure 3).

Contrary to the microsatellite data, the SNP chip data from the same study ^3^ groups the Exeter population apart from all other CA populations including those in central CA, consistent with our genome-wide SNP data. Unfortunately, their SNP chip data clustering results did not include samples from the southeast USA preventing direct comparison with their SNP clustering result. This, however, could support the view that the Exeter population, introduced in 2014 is distinct from all other CA populations and that it was introduced independently, rather than resulting from local spread of *Ae. aegypti* within CA.

Pless, et al. ^3^ PCA analyses of the SNP chip data separated Clovis (GC3) from the GC2 cluster with some overlap. The larger number of SNPs used in our analysis (>2.9 million biallelic SNPs compared to 15,698 SNPs) may have increased the resolution, allowing us to confidently separate the two. Our data together with previous reports strongly support multiple introductions of *Ae. aegypti* in California. The most likely scenario includes four independent introductions: (i) Clovis area; probably in 2013 (ii) Madera area; probably in 2013 (iii) southern CA, probably in 2014 (iv) Exeter, probably in 2014 introduced from someplace in the southeast USA like Florida. The years are based on reports from the California vector control districts. This scenario is also in line with most of the results published based on microsatellites and SNP chip data ^3^. From our data the origin of the introductions remains dubious with only the Exeter population showing signs of presumable derivation from the southeast USA. The degree of genetic differentiation found in the Clovis population between the years 2013 and 2016 (Figure 4D) indicates the population is undergoing rapid changes in its genome, potentially reflecting local adaptation, or, less likely, drift. The only other longitudinal investigation of a CA population of *Ae. aegypti* that we are aware of compares genotypes of samples from 2013 and 2015 from Madera, detecting almost no change within these two years ^3^. Further investigation on genic features showing significant differences between the two time points may shed light on the genes involved in local adaptation at Clovis and the particular circumstances that drove it.

The geographic origin of CA *Ae. aegypti* populations and the means by which they were introduced remains unclear. Perhaps the most interesting open question is what conditions facilitated multiple introductions? Answering these questions is beyond the scope of this study and requires additional data. Investigating samples from different origins using the same NGS platform may provide a clearer description of *Ae. aegypti* invasion history in CA. In addition, investigation describing genomic changes over time may provide information on local adaptation and potentially will be useful for the control of the species in California.

## Acknowledgments

We thank Orange County Mosquito and Vector Control District team (Mr. Michael Hearst, District Manager), Community Health Division of the Department of Environmental Health (Ms. Rebecca Lafreniere, Chief and Ms. Elizabeth Pozzebon, Director), Greater Los Angeles Mosquito and Vector Control District (Dr. Susan Kluh), and Dr. Christopher Barker (UC Davis) for providing specimens. We thank Youki Yamasaki, Allison Chang, Parker Houston, Allison Weakley, Kendra Person, Hans Gripkey for assisting DNA extraction and library preparations for this study. Special thanks to Dr. Bradley Main for his comments to the manuscript. This project was funded by the UC Davis Bridge Funding Program, School of Veterinary Medicine Vector-borne Disease Pilot Grant Program, DARPA Safe Gene Program and the CDC Pacific Southwest Regional Center of Excellence grant. We thank personnel from Consolidated Mosquito Abatement District and Delta, Greater LA County and San Mateo County Vector Control Districts and Fresno, Madera County and Orange County Mosquito and Vector Control Districts and San Diego County Dept. of Environmental Health, Vector Control for collecting samples used in this study. We also thank Dr. Leo Braack (University of Pretoria, South Africa) for assisting AJC in collection of South African samples and Dr. Danny Governer and SANParks for permitting collection from Shingwedzi in the Kruger National Park.

## Supplementary materials

**Table S1.** Sample metadata including geographic coordinates, collection date and genome sequencing related metric.

**Table S2.** Pairwise F_ST_ values based on nuclear genome SNP data.

**Figure S1.** Genome-wide comparison of *Ae. aegypti* populations comparing GC2 vs GC3, GC2 vs GC4 and GC3 vs GC4.

## Co-Author Contact Information

Hanno Schmidt, Vector Genetics Laboratory, Department of Pathology, Microbiology and Immunology, School of Veterinary Medicine, University of California - Davis, CA 95616, USA; Travis C. Collier, Daniel K. Inouye US Pacific Basin Agricultural Research Center (PBARC), United States Department of Agriculture, Agricultural Research Service, Hilo, Hawaii, USA; William R. Conner, Department of Entomology and Nematology, College of Agricultural and Environmental Sciences, University of California - Davis, CA 95616, USA; Mark J. Hanemaaijer, Vector Genetics Laboratory, Department of Pathology, Microbiology and Immunology, School of Veterinary Medicine, University of California - Davis, CA 95616, USA; Montgomery Slatkin, Department of Integrative Biology, University of California - Berkeley, CA 94720, USA; John M. Marshall, School of Public Health, University of California - Berkeley, CA 94720, USA; Joanna C. Chiu, Department of Entomology and Nematology, College of Agricultural and Environmental Sciences, University of California - Davis, CA 95616, USA; Chelsea T. Smartt, Florida Medical Entomology Laboratory, Institute of Food and Agricultural Sciences, University of Florida, Vero Beach, FL 32962, USA; Gregory C. Lanzaro, Vector Genetics Laboratory, Department of Pathology, Microbiology and Immunology, School of Veterinary Medicine, University of California - Davis, CA 95616, USA; F. Steve Mulligan, Consolidated Mosquito Abatement District, Selma, CA 93648, USA; Anthony J. Cornel, Mosquito Control Research Laboratory, Kearney Agricultural Center, Department of Entomology and Nematology, University of California - Davis, CA 95616, USA.

## References

1. Honorio NA, Silva Wda C, Leite PJ, Goncalves JM, Lounibos LP, Lourenco-de-Oliveira R, 2003. Dispersal of Aedes aegypti and Aedes albopictus (Diptera: Culicidae) in an urban endemic dengue area in the State of Rio de Janeiro, Brazil. Mem Inst Oswaldo Cruz 98: 191–8.

2. Harrington LC, Scott TW, Lerdthusnee K, Coleman RC, Costero A, Clark GG, Jones JJ, Kitthawee S, Kittayapong P, Sithiprasasna R, Edman JD, 2005. Dispersal of the dengue vector Aedes aegypti within and between rural communities. Am J Trop Med Hyg 72: 209–20.

3. Pless E, Gloria-Soria A, Evans BR, Kramer V, Bolling BG, Tabachnick WJ, Powell JR, 2017. Multiple introductions of the dengue vector, Aedes aegypti, into California. PLOS Negl Trop Dis 11: e0005718.

4. Gloria-Soria A, Brown JE, Kramer V, Hardstone Yoshimizu M, Powell JR, 2014. Origin of the dengue fever mosquito, Aedes aegypti, in California. PLOS Negl Trop Dis 8: e3029.

5. Cornel AJ, Holeman J, Nieman CC, Lee Y, Smith C, Amorino M, Brisco KK, Barrera R, Lanzaro GC, Mulligan Iii FS, 2016. Surveillance, insecticide resistance and control of an invasive Aedes aegypti (Diptera: Culicidae) population in California. F1000Res 5: 194.

6. Jewell D, Grodhaus G, 1984. Laird M, ed. Commerce and the Spread of Pests and Disease Vectors.. New York: Praeger Publishers, 103–107.

7. Main BJ, Lee Y, Collier TC, Norris LC, Brisco K, Fofana A, Cornel AJ, Lanzaro GC, 2015. Complex genome evolution in Anopheles coluzzii associated with increased insecticide usage in Mali. Mol Ecol 24: 5145–5157.

8. Main BJ, Lee Y, Ferguson HM, Kreppel KS, Kihonda A, Govella NJ, Collier TC, Cornel AJ, Eskin E, Kang EY, Nieman CC, Weakley AM, Lanzaro GC, 2016. The Genetic Basis of Host Preference and Resting Behavior in the Major African Malaria Vector, Anopheles arabiensis. PLOS Genet 12: e1006303.

9. Vicente JL, Clarkson CS, Caputo B, Gomes B, Pombi M, Sousa CA, Antao T, Dinis J, Botta G, Mancini E, Petrarca V, Mead D, Drury E, Stalker J, Miles A, Kwiatkowski DP, Donnelly MJ, Rodrigues A, Torre AD, Weetman D, Pinto J, 2017. Massive introgression drives species radiation at the range limit of Anopheles gambiae. Sci Rep 7: 46451.

10. Norris LC, Main BJ, Lee Y, Collier TC, Fofana A, Cornel AJ, Lanzaro GC, 2015. Adaptive introgression in an African malaria mosquito coincident with the increased usage of insecticide-treated bed nets. Proc Natl Acad Sci U S A 112: 815–20.

11. Hanemaaijer MJ, Collier TC, Chang A, Shott CC, Houston PD, Schmidt H, Main BJ, Cornel AJ, Lee Y, Lanzaro GC, 2018. The fate of genes that cross species boundaries after a major hybridization event in a natural mosquito population Mol Ecol.

12. Nieman CC, Yamasaki Y, Collier TC, Lee Y, 2015. A DNA extraction protocol for improved DNA yield from individual mosquitoes. F1000Res 4: 1314.

13. Yamasaki YK, Nieman CC, Chang AN, Collier TC, Main BJ, Lee Y, 2016. Improved tools for genomic DNA library construction of small insects. F1000Res, 211.

14. Bolger AM, Lohse M, Usadel B, 2014. Trimmomatic: a flexible trimmer for Illumina sequence data. Bioinformatics.

15. Matthews BJ, Dudchenko O, Kingan S, Koren S, Antoshechkin I, Crawford JE, Glassford WJ, Herre M, Redmond SN, Rose NH, Weedall GD, Wu Y, Batra SS, Brito-Sierra CA, Buckingham SD, Campbell CL, Chan S, Cox E, Evans BR, Fansiri T, Filipovic I, Fontaine A, Gloria-Soria A, Hall R, Joardar VS, Jones AK, Kay RGG, Kodali V, Lee J, Lycett GJ, Mitchell SN, Muehling J, Murphy MR, Omer A, Partridge FA, Peluso P, Aiden AP, Ramasamy V, Rasic G, Roy S, Saavedra-Rodriguez K, Sharan S, Sharma A, Smith M, Turner J, Weakley AM, Zhao Z, Akbari OS, Black WC, Cao H, Darby AC, Hill C, Johnston JS, Murphy TD, Raikhel AS, Sattelle DB, Sharakhov IV, White BJ, Zhao L, Aiden EL, Mann RS, Lambrechts L, Powell JR, Sharakhova MV, Tu Z, Robertson HM, McBride CS, Hastie AR, Korlach J, Neafsey DE, Phillippy AM, Vosshall LB, 2017. Improved Aedes aegypti mosquito reference genome assembly enables biological discovery and vector control. bioRxiv.

16. Li H, 2013. Aligning sequence reads, clone sequences and assembly contigs with BWA-MEM: Cornell University Library, arXiv:1303.3997v2.

17. Okonechnikov K, Conesa A, Garcia-Alcalde F, 2016. Qualimap 2: advanced multi-sample quality control for high-throughput sequencing data. Bioinformatics 32: 292–4.

18. Garrison E, Marth G, 2012. Haplotype-based variant detection from short-read sequencing. arXiv preprint.

19. Crawford JE, Lazzaro BP, 2012. Assessing the accuracy and power of population genetic inference from low-pass next-generation sequencing data. Front Genet 3: 66.

20. Felsenstein J, 1989. PHYLIP - Phylogeny Inference Package (Version 3.2). Cladistics 5: 164–166.

21. Hudson RR, Slatkin M, Maddison WP, 1992. Estimation of levels of gene flow from DNA sequence data. Genetics 132: 583–9.

22. Miles A, Harding N, 2016. scikit-allel - Explore and analyse genetic variation. https://github.com/cggh/scikit-allel: GitHub.

23. Hlaing T, Tun-Lin W, Somboon P, Socheat D, Setha T, Min S, Chang MS, Walton C, 2009. Mitochondrial pseudogenes in the nuclear genome of Aedes aegypti mosquitoes: implications for past and future population genetic studies. BMC Genet 10: 11.

24. Schmidt H, Hanemaaijer MJ, Cornel AJ, Lanzaro GC, Braack L, Lee Y, 2018. Complete mitogenome sequence of Aedes (Stegomyia) aegypti derived from field isolates from California and South Africa. Mitochondrial DNA Part B.

25. Danecek P, Auton A, Abecasis G, Albers CA, Banks E, DePristo MA, Handsaker RE, Lunter G, Marth GT, Sherry ST, McVean G, Durbin R, Genomes Project Analysis G, 2011. The variant call format and VCFtools. Bioinformatics 27: 2156–8.

26. Kumar S, Stecher G, Tamura K, 2016. MEGA7: Molecular Evolutionary Genetics Analysis Version 7.0 for Bigger Datasets. Mol Biol Evol 33: 1870–4.

27. Behura SK, Lobo NF, Haas B, deBruyn B, Lovin DD, Shumway MF, Puiu D, Romero-Severson J, Nene V, Severson DW, 2011. Complete sequences of mitochondria genomes of Aedes aegypti and Culex quinquefasciatus and comparative analysis of mitochondrial DNA fragments inserted in the nuclear genomes. Insect Biochem Mol Biol 41: 770–7.

28. Lee Y, Collier TC, Taylor CE, Cornel AJ, Lanzaro GC, 2017. UC Davis PopI OpenProjects-AedesGenomes. Available at: https://popi.ucdavis.edu/PopulationData/OpenProjects/AedesGenomes/. Accessed 2018.

29. Nazareno AG, Bemmels JB, Dick CW, Lohmann LG, 2017. Minimum sample sizes for population genomics: an empirical study from an Amazonian plant species. Mol Ecol Resour 17: 1136–1147.

30. Moore M, Sylla M, Goss L, Burugu MW, Sang R, Kamau LW, Kenya EU, Bosio C, Munoz Mde L, Sharakova M, Black WC, 2013. Dual African origins of global Aedes aegypti s.l. populations revealed by mitochondrial DNA. PLOS Negl Trop Dis 7: e2175.

31. Damal K, Murrell EG, Juliano SA, Conn JE, Loew SS, 2013. Phylogeography of Aedes aegypti (yellow fever mosquito) in South Florida: mtDNA evidence for human-aided dispersal. Am J Trop Med Hyg 89: 482–8.

32. Lynch M, 2007. The origins of genome architecture. Sunderland, Mass.: Sinauer Associates.

33. Havird JC, Sloan DB, 2016. The Roles of Mutation, Selection, and Expression in Determining Relative Rates of Evolution in Mitochondrial versus Nuclear Genomes. Mol Biol Evol 33: 3042– 3053.

34. Molnar RI, Bartelmes G, Dinkelacker I, Witte H, Sommer RJ, 2011. Mutation rates and intraspecific divergence of the mitochondrial genome of Pristionchus pacificus. Mol Biol Evol 28: 2317–26.

35. Marsden CD, Lee Y, Kreppel K, Weakley A, Cornel A, Ferguson HM, Eskin E, Lanzaro GC, 2014. Diversity, differentiation, and linkage disequilibrium: prospects for association mapping in the malaria vector Anopheles arabiensis. G3 (Bethesda) 4: 121–31.

36. Evans BR, Gloria-Soria A, Hou L, McBride C, Bonizzoni M, Zhao H, Powell JR, 2015. A Multipurpose High Throughput SNP Chip for the Dengue and Yellow Fever Mosquito, Aedes aegypti. G3 (Bethesda).

37. Nene V, Wortman JR, Lawson D, Haas B, Kodira C, Tu ZJ, Loftus B, Xi Z, Megy K, Grabherr M, Ren Q, Zdobnov EM, Lobo NF, Campbell KS, Brown SE, Bonaldo MF, Zhu J, Sinkins SP, Hogenkamp DG, Amedeo P, Arensburger P, Atkinson PW, Bidwell S, Biedler J, Birney E, Bruggner RV, Costas J, Coy MR, Crabtree J, Crawford M, Debruyn B, Decaprio D, Eiglmeier K, Eisenstadt E, El-Dorry H, Gelbart WM, Gomes SL, Hammond M, Hannick LI, Hogan JR, Holmes MH, Jaffe D, Johnston JS, Kennedy RC, Koo H, Kravitz S, Kriventseva EV, Kulp D, Labutti K, Lee E, Li S, Lovin DD, Mao C, Mauceli E, Menck CF, Miller JR, Montgomery P, Mori A, Nascimento AL, Naveira HF, Nusbaum C, O’Leary S, Orvis J, Pertea M, Quesneville H, Reidenbach KR, Rogers YH, Roth CW, Schneider JR, Schatz M, Shumway M, Stanke M, Stinson EO, Tubio JM, Vanzee JP, Verjovski-Almeida S, Werner D, White O, Wyder S, Zeng Q, Zhao Q, Zhao Y, Hill CA, Raikhel AS, Soares MB, Knudson DL, Lee NH, Galagan J, Salzberg SL, Paulsen IT, Dimopoulos G, Collins FH, Birren B, Fraser-Liggett CM, Severson DW, 2007. Genome sequence of Aedes aegypti, a major arbovirus vector. Science 316: 1718–23.

38. CalSurv, 2007. California Surveilliance Gateway Maps: California Vectorborne Disease Surveillance System.

39. Patterson T, 2008. CleanTOPO2: Edited SRTM30 Plus World Elevation Data. Online (author communicated that his data is published in public domain and free to use)..

